# A critique of common methods for comparing the scaling of vertical force production in flying insects

**DOI:** 10.1101/2022.02.11.480158

**Authors:** Nicholas P. Burnett, Emily Keliher, Stacey A. Combes

**Affiliations:** Department of Neurobiology, Physiology, and Behavior, University of California – Davis, Davis, CA 95616

## Abstract

Maximum vertical force production (F_vert_) is an integral measure of flight performance that generally scales with size. Numerous methods of measuring F_vert_ and body size exist, but few studies have compared how these methods affect the conclusions of scaling analyses. We compared two common techniques for measuring F_vert_ in bumblebees (*Bombus impatiens*) and mason bees (*Osmia lignaria*), and examined F_vert_ scaling using five size metrics. F_vert_ results were similar with incremental or asymptotic load-lifting, but scaling analyses were sensitive to the size metric used. Analyses based on some size metrics indicated similar scaling exponents and coefficients between species, whereas other metrics indicated different coefficients. Furthermore, F_vert_ showed isometry with body lengths and fed and starved masses, but negative allometry with dry mass. We conclude that F_vert_ can be measured using either incremental or asymptotic loading but choosing a size metric for scaling studies requires careful consideration.

## Introduction

Maximum vertical force production (F_vert_) is an integral component of flight performance, and has been examined across a diversity of volant taxa (Marden, 1987). To maintain flight altitude, animals must produce vertical forces that match their body weight (mass*gravitational acceleration), and more elaborate flight behaviors require additional force production. For instance, animals that produce vertical forces exceeding their body weight can engage in vertical acceleration (e.g., evasive flight maneuvers) or load-carrying (e.g., transporting food or nesting materials) (Buchwald and Dudley, 2010; Marden, 1987; Wolf and Schmid-Hempel, 1989). F_vert_ generally scales isometrically with flight muscle mass and body size in closely related taxa (Marden, 1987), but variation in F_vert_ scaling can exist between distantly related species due to differences in musculature, morphology, behavior, or kinematics (Chai et al., 1997; Dillon and Dudley, 2004; Marden, 1987). Although previous studies have compared techniques for measuring F_vert_ or quantifying body size in scaling analysis (Buchwald and Dudley, 2010; Cane, 1987), these assessments explored only one of the variables involved (i.e., either F_vert_ or body size). Thus, the full range of effects that different methodologies can have on the outcome of interspecific scaling studies of vertical force production remains unknown.

### Measuring vertical force production

The two simplest methods of measuring F_vert_, incremental and asymptotic load-lifting, involve challenging animals to sustain flight with the heaviest added load that they can carry (see *Results & Discussion* for a discussion of alternative methods). In the incremental method, weights are attached to an animal and the animal is prompted to fly. After each successful flight, additional weights are added. This process is repeated until the animal can no longer fly, and the maximum load (body + added weights) reached before failure defines the animal’s F_vert_. The incremental method has been used on bats, birds, and insects (Marden, 1987), and on the bumblebee *Bombus impatiens* (Buchwald and Dudley, 2010). In the asymptotic method, a beaded string (i.e., small masses attached to a string at fixed intervals) is attached to an animal and the animal is prompted to fly vertically. As the animal takes off and increases altitude, it lifts more of the beaded string until it is unable to lift any additional mass; the weight of the animal’s body plus the beads and string lifted indicates the animal’s F_vert_. This method has been used on hummingbirds (Altshuler and Dudley, 2003; Chai et al., 1997), orchid bees (Dillon and Dudley, 2004), and *B. impatiens* (Buchwald and Dudley, 2010; Mountcastle and Combes, 2013).

The asymptotic method is advantageous because F_vert_ is measured in a single flight trial, whereas the incremental method requires numerous flights. However, the asymptotic method may be problematic for species with erratic, non-vertical flight behaviors (Su et al., 2020), and both methods can be difficult in species that will not tolerate being handled or having a mass attached to their body (Altshuler and Dudley, 2003). Comparisons of these methods have suggested that the incremental method may underestimate F_vert_ (Buchwald and Dudley, 2010), but this assessment has not been replicated or tested in additional species. Assessing the validity of these widespread methods of measuring F_vert_ is necessary to facilitate comparative studies of species exhibiting flight behaviors that may preclude one of the methods.

### Scaling performance by size

All flying animals must produce, at minimum, enough force to support their own body weight, so F_vert_ generally increases with body size. Previous work has shown that F_vert_ often increases isometrically with body size (Buchwald and Dudley, 2010; Marden, 1987; Marden, 1990), but with some exceptions (Dillon and Dudley, 2004). In bees (Apoidea), body size is commonly quantified using either a length measure (e.g., wing length, intertegular (IT) span) or a mass measure (e.g., fed, starved, or dry body mass). Are these metrics interchangeable for scaling analyses of flight performance? IT span (the distance between the tegulae at the wing bases) and wing length are morphological features that can be measured directly with calipers or through photographs, and are proportional to body mass in closely related taxa (Cane, 1987; Dillon and Dudley, 2004). Fed body mass is the body mass measured at some point before or after a flight trial; this measure can introduce variability because each bee may be carrying a different volume of nectar when selected for a flight trial (Marden, 1987). Starved (or empty) body mass is the body mass of a bee without any stored nectar, and thus, this measure represents the baseline body mass that the bee must lift to fly; it can be obtained by measuring body mass after squeezing a bee to cause regurgitation of nectar from the bee’s honey sac, or crop (Buchwald and Dudley, 2010). However, this technique can introduce error because not all nectar is stored in the crop – up to 10% can be retained in the bee’s midgut after regurgitation of the crop’s contents (Gary and Lorenzen, 1976). Alternatively, empty body mass can be obtained by weighing the bee after it is starved over some time period (e.g., 24 hours) to allow the bee to metabolize all of the nectar in its body, without allowing the bee to desiccate or die (Combes et al., 2020). Dry body mass is the body mass after desiccating a dead bee in an oven until it stops losing weight (Cane, 1987; Helm et al., 2021); this method can introduce error because solutes within nectar (or other fluids in the body) may remain in the bee after desiccation (especially if the bee was not starved beforehand), adding to the dry mass. Although IT span, wing length, and fed, starved, and dry body masses are among the simplest and most widespread body size measurements used in insect flight studies, the variability introduced by each of these metrics has not been compared across species in the context of flight performance.

### Study system

Here, we compare two simple methodologies for quantifying F_vert_, by performing both measurements on individual females of two bee species, the eastern bumblebee *Bombus impatiens* and the mason bee *Osmia lignaria*. We then test whether interspecific comparisons of flight performance, controlled for body size, depend on the body size metric used in the analyses. These species are in the superfamily Apoidea but differ in body size (most *O. lignaria* females are smaller than *B. impatiens* workers), morphology, and life history (*O. lignaria* are solitary and B. *impatiens* are primitively eusocial). *Bombus impatiens* is an established model organism for flight biomechanics studies (Buchwald and Dudley, 2010; Combes et al., 2020; Mountcastle and Combes, 2013), and *O. lignaria* is an emerging model for studies of flight biomechanics, reproductive physiology, and landscape ecology (Bosch and Kemp, 2000; Helm et al., 2021; Kemp et al., 2004; Vicens and Bosch, 2000). Both species are also sold commercially for use in crop pollination, as an alternative to honeybees. Thus, these species represent taxa that are not closely related, but may be inadvertently grouped together in broad analyses of flight performance across taxa.

## Materials and methods

Cocoons of adult-wintering *Osmia lignaria* were purchased from a commercial supplier (Foothill Bee Ranch, Auburn, CA, USA) and maintained at 4°C. Individuals were moved to a flight cage for emergence, as needed for experiments. A mature colony of *Bombus impatiens* was purchased from a commercial supplier (Koppert Biological Systems, Romulus, MI, USA) and maintained in a separate flight cage. Individuals in each cage were fed sucrose solution *ad libitum*, with fresh pollen weekly. All flight cages and experimental areas were held at 22-25°C. Active females (n = 25 for *O. lignaria*, n = 28 for *B. impatiens)* of each species were selected randomly for flight trials.

### Flight performance

F_vert_ was measured on each individual using both the incremental and asymptotic methods, to allow for direct comparison. The order of the methods was alternated between individuals, with both tests performed during the same day, and testing methodology generally followed the descriptions by Buchwald and Dudley (2010) and Mountcastle and Combes (2013). Briefly, bees were cold-anesthetized at 4°C and a polyester thread was tied around the petiole of each individual (Mountcastle and Combes, 2013), near their center of lift (Buchwald and Dudley, 2010; Dudley and Ellington, 1990; Ellington, 1984a), leaving a free end of thread approximately 6 cm long. Once tied, bees were allowed to recover at room temperature for 10-20 minutes before any flight trials.

#### Incremental method

Individual beads (either 0.0250 or 0.0050 g in mass) were tied to the free end of thread around a bee’s petiole. Prior to each flight trial, the mass of the bee, string, and beads were recorded. The bee was released into a flight arena and prompted to fly, using agitation with forceps if necessary (Fig. 1a). If the bee took off and sustained flight, additional beads were added, mass was recorded, and the flight trial was repeated until the bee was unable to fly with the weight applied. The maximum mass lifted by the bee was multiplied by gravitational acceleration to calculate the bee’s F_vert_.

**Figure 1.**
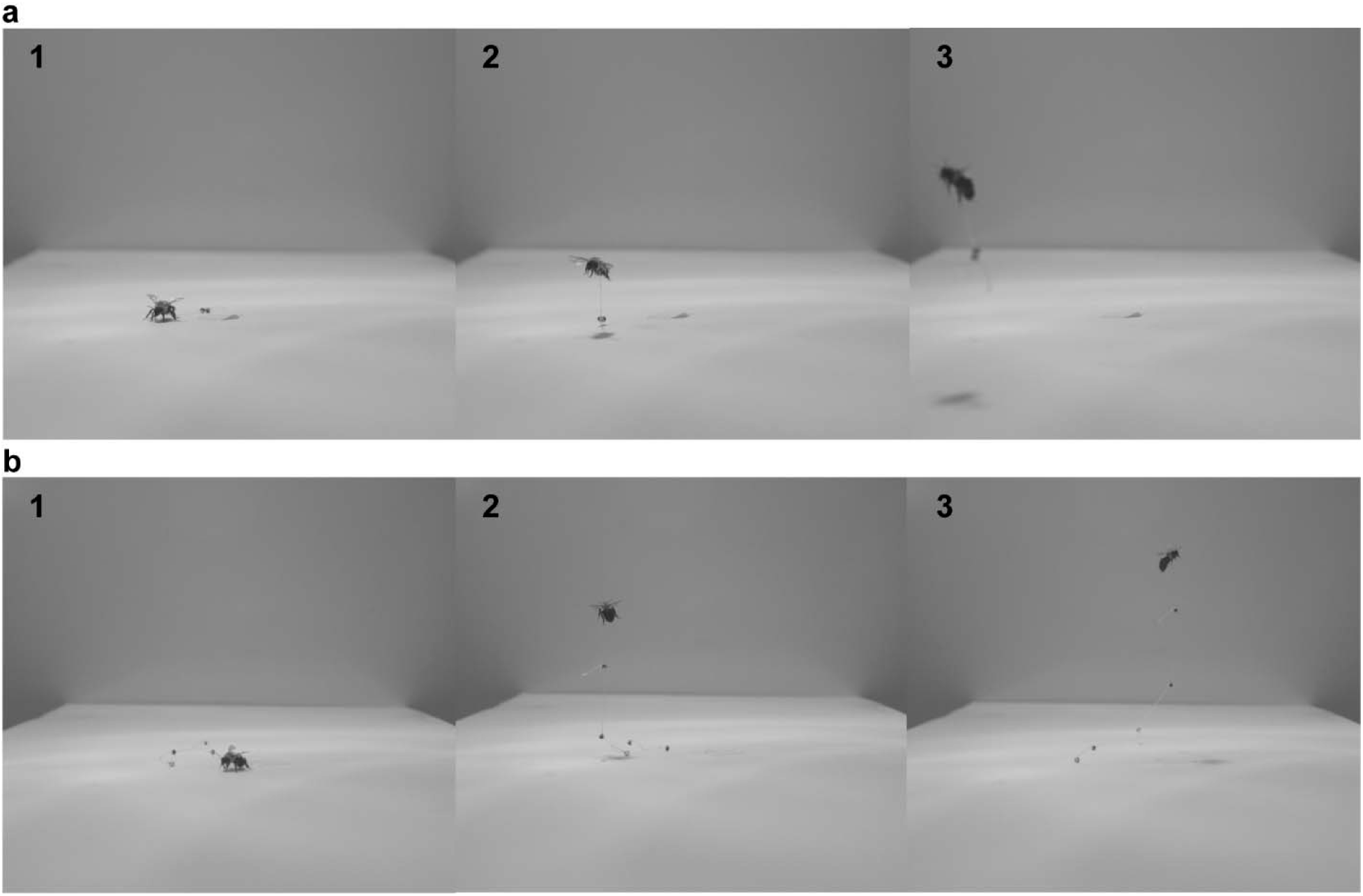
Examples of the (a) incremental and (b) asymptotic methods for measuring maximum vertical force production. Each example shows a three-photograph sequence of a single flight attempt, using a female *Bombus impatiens*.

#### Asymptotic method

Beads (either 0.0250 or 0.005 g in mass) were attached to a polyester string, approximately 30 cm in length, at intervals of 2 cm. Based on preliminary trials, the strings used in *O. lignaria* flight tests had beads with mass = 0.0050 g, and the strings used in *B. impatiens* flight tests had beads with mass = 0.0250 g. Before and after each flight trial, the mass of the bee, along with the 6-cm thread tied around its petiole, was measured. During a flight trial, the bee was tied to the beaded string and prompted to fly, using agitation with forceps if necessary (Fig. 1b). Flights were recorded with a video camera at 30-60 frames per second, and the maximum number of beads lifted during each sustained vertical flight was counted. Up to five successful flight trials were recorded per individual. F_vert_ was calculated as the sum of bee mass (averaged between the pre- and post-flight mass) and the lifted mass of the beaded string, multiplied by gravitational acceleration. The lifted mass of the beaded string was calculated as the maximum number of beads lifted during the flights, multiplied by the average mass per bead (total mass of string and bead, divided by the number of beads on the string).

In both incremental and asymptotic methods, we consider the maximum lifted mass to be the observed maximum lifted mass, following Mountcastle and Combes (2013). However, other studies have considered the maximum lifted mass to be the mean between the observed maximum lifted mass and the next-highest mass that the bee was unable to lift (Buchwald and Dudley, 2010; Marden, 1987). While this variation in methodology can impact comparisons of data between studies, it does not affect the conclusions of the present study because the same approach was used for all trials.

### Body size

After all flight trials using both methods were completed for each bee, the string was removed from the petiole and body mass was measured to the nearest 0.0001 g with a digital balance (providing the fed mass). The bee was placed in a separate dish with only a wet paper towel and left for 24 h at room temperature to consume any nectar remaining in its body. After 24 h, body mass was measured again (providing the starved mass), and the bee was placed in a freezer until all experiments were completed.

Once all flight tests were completed, we removed bees from the freezer, photographed them, and measured their intertegular (IT) span and forewing length (hereafter wing length) to the nearest 0.01 mm using ImageJ (v 1.53f51) (Schneider et al., 2012). Following these geometric measurements, bees were placed in a drying oven at 45°C, following Cane (1987), and dried for several days until all bees were no longer losing mass (providing the dry mass).

### Statistical analysis

Within each species, we compared F_vert_ measurements between the incremental and asymptotic methods using paired *t-*tests (paired by individual).

We next compared F_vert_-size scaling between the species using the five body size metrics described above. F_vert_ and size data for each species can be represented by the power function *Y* = β*X^α^*, where *Y* is F_vert_, *β* is a scaling coefficient, *X*is body size, and *α* is a scaling exponent. This power function can also be expressed in a logarithmic form: log_10_ *Y* = log_10_*β* + *α*log_10_ *X* Here, β is the *Y*-intercept of the log_10_-transformed model and α is the slope of the log_10_-transformed model (Vogel, 2013).

We log_10_-transformed all data to conduct an ANCOVA (analysis of covariance) scaling analysis for each body size metric, using F_vert_ as the dependent variable, species as the independent variable, and body size as the covariate. We first tested for a statistical interaction between species and body size (i.e., different scaling exponents between species); if none was found, we then tested for a statistical effect of species across body size (i.e., different scaling coefficients between species). Last, we tested whether F_vert_ scaled isometrically (i.e., a scaling exponent = 1 for body masses, 3 for body lengths) with each body size metric, using Wald tests. All analyses were done in R Statistical Software (R Core Team, 2020).

## Results and Discussion

### Size metrics affect the outcome of scaling analyses, but F_vert_ methods do not

F_vert_-size scaling depended strongly on the size metric used, but not on the method of measuring F_vert_. The incremental and asymptotic methods for quantifying F_vert_ produced similar results within each species (Fig. 2). For *Osmia lignaria,* the methods differed by 3.0 ± 10.4% (mean ± SD), calculated as asymptotic - incremental, divided by the average of the two methods (paired *t*-test, *t* = −1.310, df = 19, *p* = 0.206). For *Bombus impatiens,* the methods differed by 1.6 ± 12.6% (*t* = 0.808, df = 20, *p* = 0.428).

**Figure 2.**
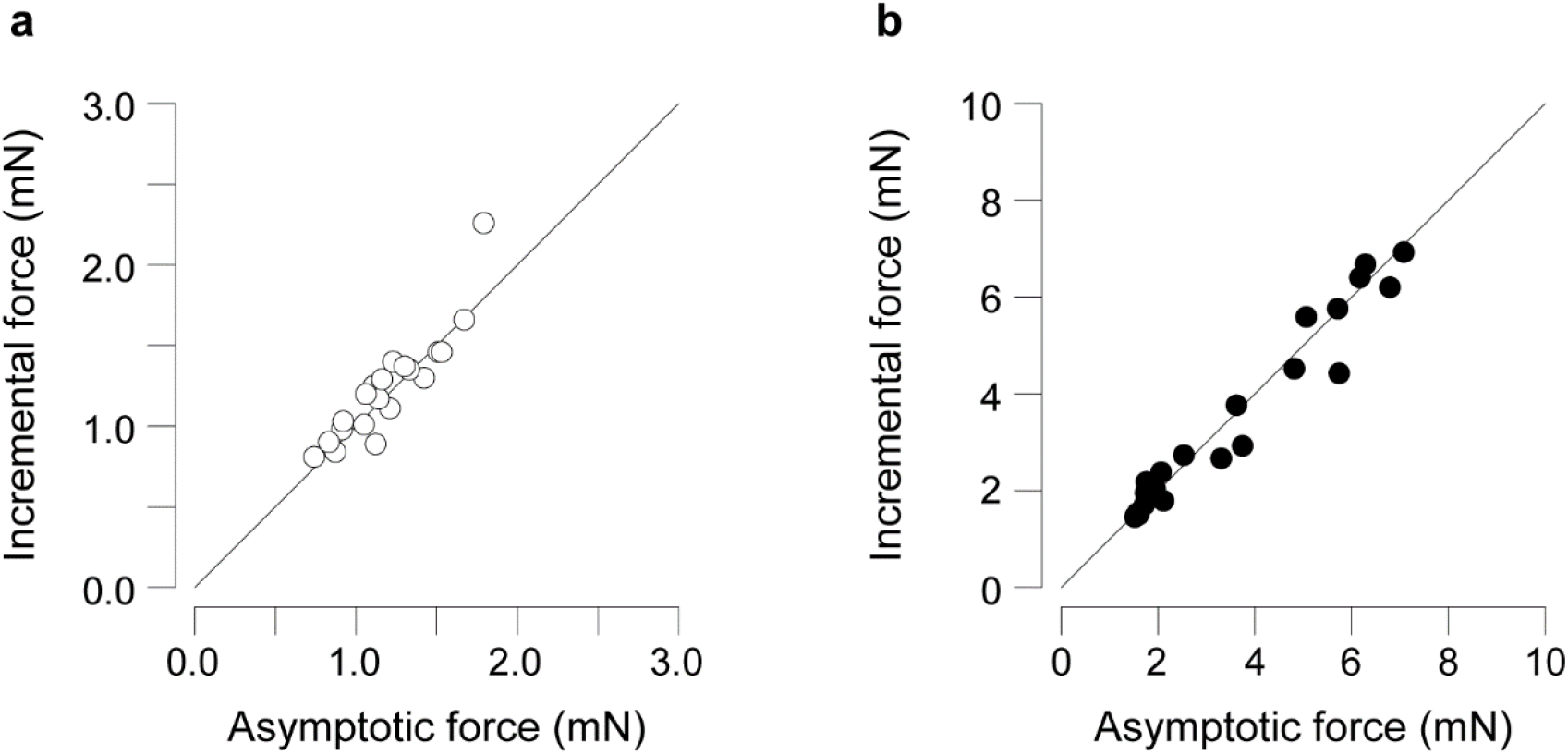
Paired measurements of vertical force production show that the incremental and asymptotic methods produce similar results. Paired F_vert_ measurements using both methods in each individual are shown for (a) *Osmia lignaria* (n = 25) and (b) *Bombus impatiens* (n = 28). Horizontal axes show F_vert_ measured with the asymptotic method and vertical axes show F_vert_ measured with the incremental method. The line in each panel shows a slope = 1. In both cases, incremental and asymptotic methods produced statistically similar results (paired *t*-tests, *p* > 0.05).

Plotting F_vert_ as a function of the five body size metrics revealed no differences in scaling exponent (i.e., slope of the log_10_-transformed data) between species (ANCOVA, *p* > 0.05). However, the scaling coefficient (i.e., intercept of the log_10_-transformed variables) differed significantly between species (*p* < 0.005) when IT span or dry mass was used as the size metric but was similar between species with the other three size metrics.

Using values from the incremental method (asymptotic results are similar), F_vert_ scaled isometrically (expected scaling exponent *α* = 3) with wing length (*α* = 2.778; F_(1,44)_ = 3.942, *p* = 0.053) and IT span (*α* = 2.711; F_(1,43)_ = 3.262, *p* = 0.079). F_vert_ also scaled isometrically (expected *α* = 1) with fed mass (*α* = 1.052; F_(1,45)_ = 2.493, *p* = 0.121) and starved mass (*α* = 0.980; F_(1,47)_ = 0.450, *p* = 0.506) (Fig. 3), but showed negative allometry with dry mass (*α* = 0.851; F_(1,44)_ = 11.414, *p* < 0.005). With IT span and dry mass, the scaling coefficient *β* was 0.187 and 0.137 lower, respectively, for *O. lignaria* than for *B. impatiens* (*p* < 0.005)

**Figure 3.**
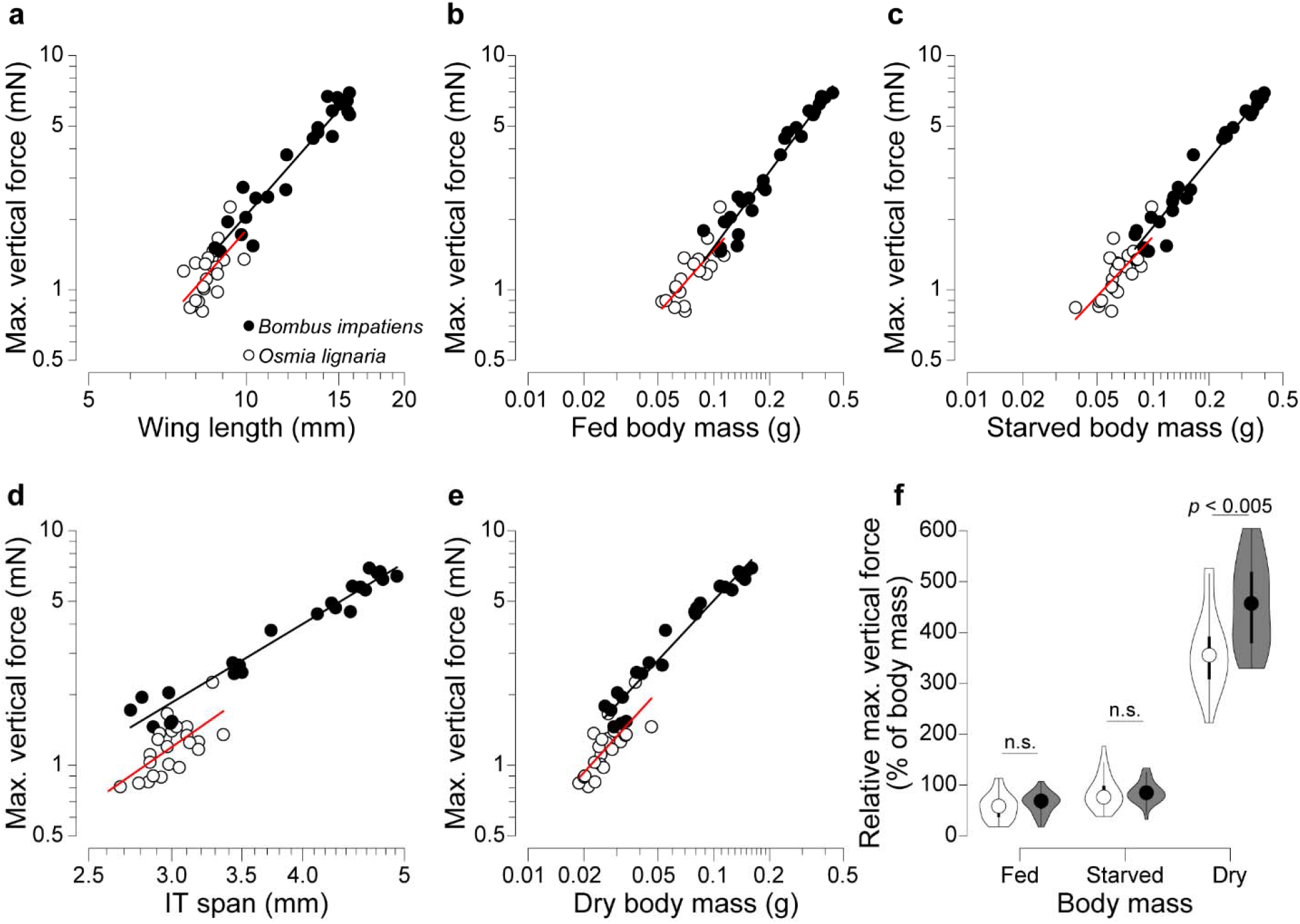
Inter-specific scaling analyses of maximum vertical force production lead to different conclusions depending on the size metric used. *Bombus impatiens* and *Osmia lignaria* display statistically similar scaling exponents and coefficients when F_vert_ is expressed as a function of (a) wing length, (b) fed body mass, or (c) starved body mass (ANCOVA, *p* > 0.05). The two species display similar scaling exponents but significantly different coefficients (i.e., *Y*-intercepts) when F_vert_ is expressed as a function of (d) IT span or (e) dry body mass (*p* < 0.005). (f) Mass-specific F_vert_ is similar in *B. impatiens* and *O. lignaria* if F_vert_ is normalized to fed or starved mass, but significantly larger in *B. impatiens* if F_vert_ is normalized to dry mass (*t*-tests, *p* < 0.05 for significance). In (f), circles show medians, bars show 25^th^ and 75^th^ percentiles, and violin plots shown the kernel density-smooth representations of the frequency distributions. White symbols represent *O. lignaria*, and black/gray symbols represent *B. impatiens*.

F_vert_ scaling was nearly identical between *O. lignaria* and *B. impatiens* when wing length, fed mass, and starved mass were used as size metrics. Thus, analyses using these metrics would suggest that *B. impatiens* produces more vertical force only because it is a larger bee than *O. lignaria,* and that both species have similar F_vert_ when normalized to their fed or starved mass (paired *t*-tests,*p* > 0.05; Fig. 3f). However, different scaling patterns emerge with IT span and dry mass, and F_vert_ normalized to dry mass differs significantly between species (*p* < 0.005). This would suggest that interspecific variation in F_vert_ is not due solely to variation in body size, but rather to some other factor such as physiology or kinematics.

### Alternative methods and considerations for future studies

Here we show that the incremental and asymptotic methods of measuring F_vert_ produce equivalent results for *B. impatiens* and *O. lignaria.* This suggests that either method can measure F_vert_ accurately, and that researchers can choose whichever method is most feasible given their study subjects’ flight behavior. However, not all previous studies comparing these methods follow the same pattern. For instance, Buchwald and Dudley (2010) found that the incremental method systematically underestimated F_vert_ in *B. impatiens*, a result that could be due to their different method of applying incremental weights (gluing rather than tying weights to bees). Estimates of F_vert_ can also depend on whether the assay involves a steady flight behavior (hovering or slow, level flight, as in the incremental and asymptotic methods) or a dynamic flight behavior (rapid accelerations). Su et al. (2020) quantified F_vert_ in a dragonfly (*Pantala flavescens*) using a dynamic flight behavior, in which the animal was loaded with weights and then dropped. The dragonfly’s rapid acceleration as it stopped its fall towards the floor and then ascended upwards (a “pull-up” response) can be used to calculate F_vert_ (mass*maximum vertical acceleration). However, these values were much higher than measurements from similar species in which sustained flight assays of F_vert_ were used (Su et al., 2020), suggesting that assays for F_vert_ based on sustained flight behavior may be broadly incompatible with assays based on dynamic flight behavior.

Another common tool for assessing maximum flight performance is the variable gas mixture method. With all else being equal, the vertical force (lift) produced by a flying animal will decrease as air density decreases (Vogel, 1996). Variable gas mixtures can be used to alter air density while maintaining constant oxygen concentration, to identify the minimum air density in which a flying animal can generate enough vertical force to support its own body weight (Dudley and Chai, 1996; Roberts et al., 2004). When filmed with high-speed cameras, measured wing kinematics and morphological details can be used to estimate the animal’s average vertical force production (Dudley, 1995), by using the hovering aerodynamic model of Ellington (1984b). Thus, variable gas mixtures can provide an informative metric of peak flight performance, but the methodology requires significantly more equipment (sealed chamber, pressurized gas canisters, high-speed cameras) and detailed kinematic analysis, and the data are not directly comparable to force measurements from the incremental and asymptotic load-lifting methods, or even the dynamic F_vert_ assays using the loaded “pull-up” response.

Like flight performance assays, body size metrics may not always be interchangeable or comparable, especially between distantly related species. For instance, IT span may be useful for comparing size within bee species, but tegulae (and thus IT span) are only found in certain insect groups. Single linear dimensions of animals may also be misleading, as three-dimensional differences in morphology between species or across ontogeny may not be captured by a linear measurement. In some past studies, the scaling of flight performance across large and diverse groups of organisms has been examined using flight muscle mass (rather than total body mass) as the size metric, because the flight muscles are the tissues that actuate the wings (Buchwald and Dudley, 2010; Dudley, 1995; Marden, 1987; Marden, 1990). However, different species – and even individuals of different sizes within a species – may require different techniques for isolating flight muscle, which could bias morphological comparisons. For instance, flight muscle in bees and other insects can be quantified via dissection or chemical digestion of the thorax, and each technique has its own unique sources of error (e.g., correctly dissecting or digesting all of the flight muscle, and *only* flight muscle) (Buchwald and Dudley, 2010; Dudley, 1995; Marden, 1987). Thus, it is imperative for researchers to confirm that the size metrics used in inter- or intraspecific comparisons of flight performance are compatible across the range of organisms studied.

